# Sleep Facilitates Pattern Separation through SK Channel-Mediated Sparse Coding

**DOI:** 10.1101/2025.10.08.681112

**Authors:** Chien-Chun Chen, Yu-Chun Huang, Antonio Ortega, Raquel Suárez-Grimalt, Erik Tedre, El-Sayed Baz, Yifan Wu, Andrew C. Lin, Sha Liu

## Abstract

The roles of sleep in priming the brain for associative learning remain unclear. Here, we report that acute sleep deprivation in *Drosophila* selectively impairs pattern separation—the ability to distinguish between similar stimuli—without affecting classical conditioning. This deficit correlates with disrupted sparse coding in the mushroom body, reflected by an increased number of active Kenyon cells and greater overlap in their odor representations. Electrophysiological analyses reveal that sleep loss enhances small conductance calcium-activated potassium (SK) channel-mediated afterhyperpolarization in GABAergic anterior paired lateral (APL) neurons, leading to reduced levels of feedback inhibition onto Kenyon cells and compromised sparse coding. Targeted knockdown of SK channels in APL neurons reduce their augmented afterhyperpolarization and rescues the pattern separation deficits caused by sleep deprivation. These findings identify a critical role for SK channels in inhibitory interneurons to enable sleep to preserve sparse and decorrelated neural representations, supporting cognitive processes such as pattern separation.

## Introduction

Sleep is essential for various cognitive functions, notably learning and memory^1–5^. While substantial research has elucidated the mechanisms by which post-learning sleep facilitates memory consolidation, the immediate effects of the sleep before learning on subsequent information processing and memory encoding remain less understood. Recent human studies have reported that acute sleep deprivation selectively impairs certain aspects of associative learning. Specifically, it disrupts pattern separation, a neural mechanism that transforms similar or overlapping input patterns into distinct representations—allowing precise discrimination of similar stimuli during associative learning^6^. However, whether such sleep-dependent deficits occur in non-human species is unclear, and the underlying neural mechanisms remain to be elucidated.

*Drosophila melanogaster* offers a valuable model to investigate these questions. Fly sleep shares many features with mammalian sleep, including its roles in learning and memory^7–11^. Moreover, the fly’s mushroom body—a central brain structure involved in learning and memory—employs sparse coding, in which each stimulus activates only a small, distinct subset of neurons^12–15^. This sparse, decorrelated activity pattern minimizes overlap between representations of similar stimuli and enhances pattern separation^16,17^. Thus, this well-characterized neural architecture, combined with the genetic and circuit-level tools available in *Drosophila*, provides a unique opportunity to dissect how sleep loss before learning impacts subsequent information processing and memory formation at behavioral, circuit, and cellular levels.

Here, we report that acute sleep deprivation in *Drosophila melanogaster* selectively impairs pattern separation during associative learning—mirroring results observed in humans. Mechanistically, this deficit stems from sleep-deprivation-driven upregulation of small conductance calcium-activated potassium (SK) potassium channel^18^ activity in the GABAergic anterior paired lateral (APL) neurons, which in turn reduces the levels of feedback inhibition and alters the sparse coding in the mushroom body. Given that sparse coding is a conserved neural coding scheme—employed in regions such as the mammalian cortex^19–21^, hippocampus^22–26^, and insect mushroom body^27,28^—and underlies pattern separation, our findings indicate a broad, evolutionarily conserved mechanism. Specifically, sleep regulates feedback inhibition in neural circuits, tuning the sparsity of representations to support cognitive processes like pattern separation.

## Results

### Acute Sleep Loss Impairs Pattern Separation in *Drosophila*

To investigate how acute sleep deprivation impacts associative learning, flies were sleep-deprived overnight for 12 hours from Zeitgeber time (ZT) 12 to ZT24 and tested for their ability to form associations between odor stimuli and electric shocks in the following morning. We utilized the individual *Drosophila* olfactory conditioner (iDOC), a single-fly adaptation of the classical T-maze paradigm recently developed in our laboratory^29^, to evaluate associative learning. iDOC enables high-throughput aversive olfactory conditioning and precise measurement of learned performance at the individual fly level. Assessing individual fly performance was critical since sleep monitoring and deprivation were conducted on a per-fly basis, allowing us to directly link sleep metrics with learning performance within the same animal.

We first assessed whether classical olfactory aversive learning remained intact after sleep loss. Flies were given a pre-training choice between an odor stimulus, 3-octanol (OCT), and clean air to measure baseline odor preference. After conditioning, where OCT was paired with electric shock, we re-tested their odor preference. Both sleep-deprived and control (normal sleep) flies learned to avoid the conditioned stimulus (OCT in this case), showing a significant drop in performance index after training (Figure S1). These results indicate that acute overnight sleep deprivation does not impair classical Pavlovian learning in *Drosophila*.

Next, we investigated whether acute sleep deprivation affects the ability to discriminate between similar odor mixtures, a behavioral readout of pattern separation. Flies were trained to associate an odor mixture of OCT and methylcyclohexanol (MCH) at 4:1 ratio (4OCT:1MCH) with electric shock and then assessed with a choice between this mixture and a similar, non-shocked odor mixture (1OCT:4MCH; Figure 1A). Flies with normal sleep successfully learned to avoid the conditioned odor mixture, indicating that they could distinguish between highly similar odor mixtures (Figure 1B). In contrast, sleep-deprived flies failed to show learned avoidance under the same conditions, suggesting impaired discrimination of similar odor stimuli (Figure 1C). Notably, three hours of recovery sleep following 12 hours sleep loss restored the ability to discriminate these similar odor mixtures (Figure 1D), implying that sleep facilitates pattern separation.

**Figure 1.**
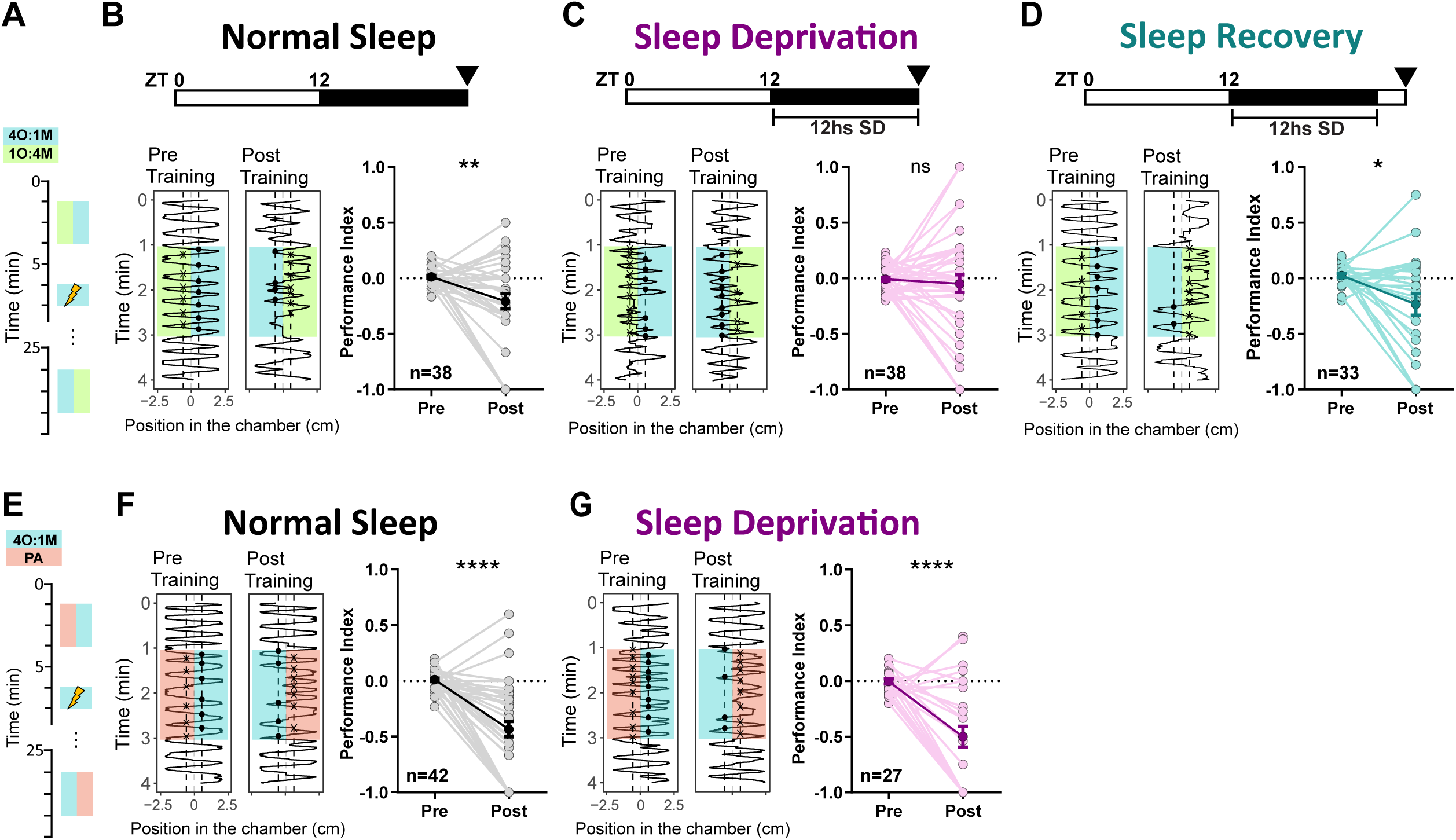
Acute sleep deprivation impaired learned discrimination of similar but not dissimilar odor pairs. **(A)** Experimental paradigm illustrating the learned odor discrimination task, comprising pre-training test, aversive conditioning (training), and post-training test phases plotted vertically against time. Colored rectangles indicate odor presentations. For similar odor discrimination, blends of OCT and MCH were used at ratios of 4:1 (4O:1M; CS^+^) and 1:4 (1O:4M; CS^−^). **(B-D)** Discrimination performance for similar odor mixtures. Upper: bars show 12:12 light-dark cycles (white: light, black: dark); Zeitgeber Time 0 (ZT0) indicates the light on; arrows indicate the timing of the learned odor discrimination task. Left: representative trajectories of fly positions in the chamber (horizontal dimension) over time (vertical dimension) before and after conditioning. Right: quantification of learning (performance index). **(B)** Normal sleep flies (n=38) successfully learned to discriminate similar odor mixtures, with the performance index significantly reduced after conditioning. **(C)** Sleep-deprived (SD) flies (12h; n=38; p= 0.6379) failed discrimination. **(D)** Sleep recovery flies (3h recovery after 12h deprivation; n=33) regained discrimination. **(E)** Experimental paradigm of the dissimilar odor mixture discrimination task, with 4O:1M as CS^+^ and PA only as CS^−^. **(F, G)** Representative trajectories and learning performance quantification are presented similarly as described above. Both **(F)** normal sleep (n=42) and **(G)** sleep deprivation groups (n=27) successfully learn to discriminate the dissimilar odors. Individual fly performance indices are shown as light-colored dots; mean values are represented by dark-colored dots. Error bars represent SEM. Statistical significance was examined by Wilcoxon matched-pairs signed rank test; ns, not significant; *P < 0.05, **P < 0.01, ***P < 0.001, ****P < 0.0001.

Importantly, this deficit was specific to similar odor discrimination. Sleep-deprived flies trained to discriminate between the 4OCT:1MCH mixture and a chemically distinct odor, propionic acid (PA), showed robust learning comparable to non-sleep-deprived control flies (Figures 1E–1G). These findings suggest that sleep loss selectively impairs the discrimination of similar odor stimuli in *Drosophila*, mirroring observations in human studies in which sleep deprivation disrupts pattern separation.^6,30^

### Sleep Maintains Sparse Coding in the Mushroom Body

In *Drosophila*, the mushroom body employs sparse and decorrelated coding of sensory information, a neural coding scheme critical for similar odor discrimination.^15^ Sparse coding ensures that only a limited subset of neurons respond to a given stimulus, and each neuron is activated by only a few stimuli^20,27^. Specifically, for any given odor, approximately 5–10% of excitatory principal neurons in the mushroom body, known as Kenyon cells, are activated, and different odors elicit responses in largely non-overlapping Kenyon cell populations^12,14,15^.

Given that acute sleep deprivation impairs pattern separation during associative olfactory learning in flies, we hypothesized that sleep loss would disrupt the sparse coding in the mushroom body. To test this, we expressed the genetically encoded calcium indicator GCaMP6f^31^ in all Kenyon cells and performed *in vivo* calcium imaging of Kenyon cell somata in response to a panel of six odors, including odors used in previous behavioral experiments (Figure 2A). Our functional imaging experiments revealed that sleep-deprived flies exhibited a greater number of Kenyon cells responding to each odor compared to control flies with normal sleep (Figure 2B), reflecting a significant reduction in population sparseness following sleep loss (Figure 2C). Furthermore, the similarity of Kenyon cell activation patterns between different odors generally increased after sleep deprivation, as measured by pairwise correlation analyses (Figure 2D). Notably, the similarity between responses to similar odor pairs (4OCT:1MCH and 1OCT:4MCH) was significantly higher in sleep-deprived flies (Figure 2E), while correlations between dissimilar odor pairs (4OCT:1MCH and PA) remained unchanged (Figure 2F). Collectively, these findings suggest that 12 hours acute sleep deprivation impairs both sparse and decorrelated representations of odors in the mushroom body, leading to less distinct neural codes.

**Figure 2.**
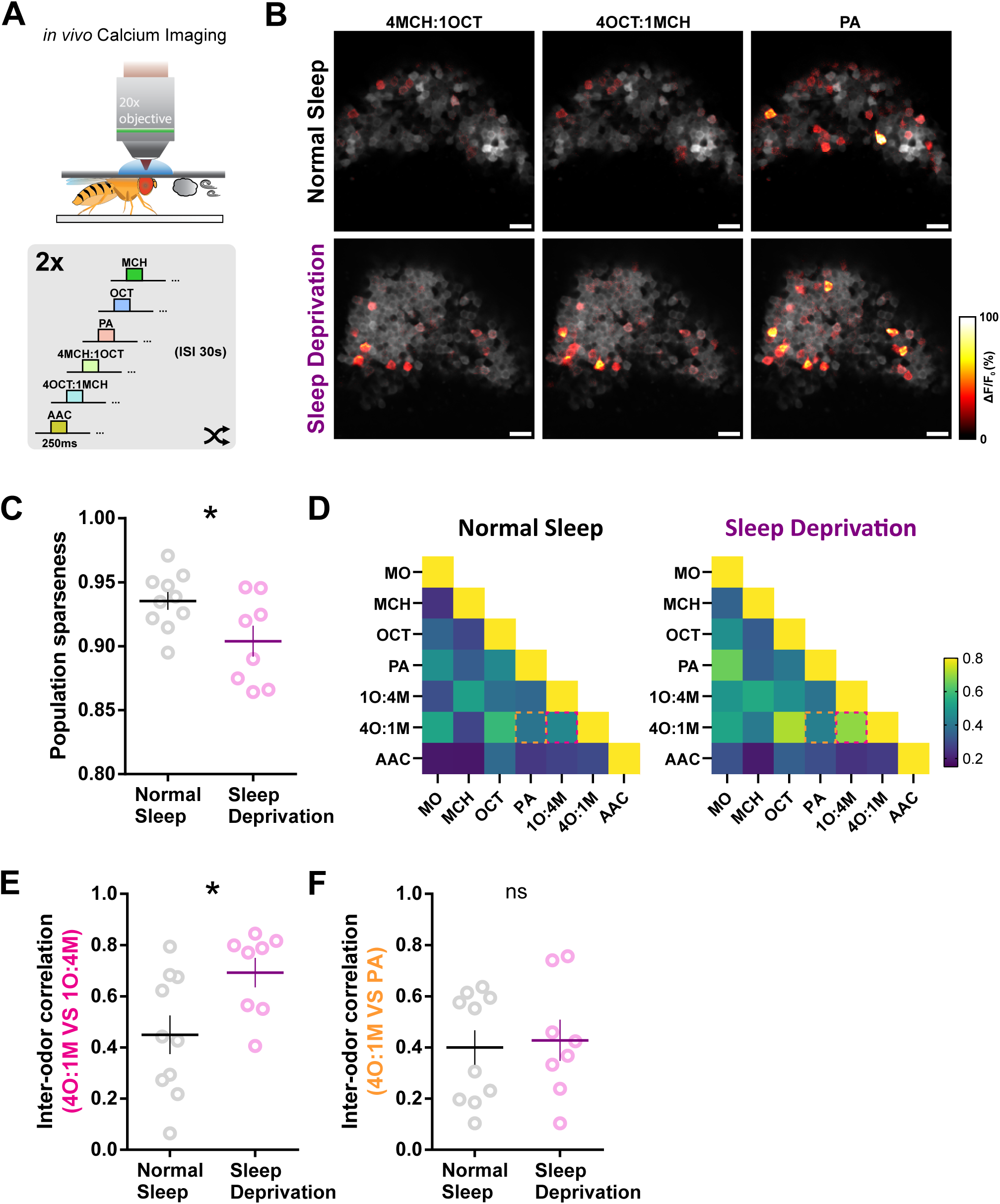
Sleep deprivation decreases population sparseness and disrupts decorrelation of olfactory representations in the mushroom body. **(A)** Top: Schematic illustrating *in vivo* calcium imaging in flies expressing GCaMP6f in all Kenyon cells using two-photon microscopy. Bottom: Experimental paradigm showing repeated application of a randomized sequence of odor stimuli to individual flies, with an inter-stimulus interval (ISI) of 30 sec. **(B)** Representative pseudo-colored calcium response maps in Kenyon cells from normal sleep (top) and sleep-deprived (bottom) flies, evoked by three odors used in Figure 1: 4O:1M, 4M:1O, and PA, overlaid on grayscale baseline fluorescence images. Scale bar, 10 µm **(C)** Quantification of population sparseness averaged across seven olfactory stimuli. Sleep deprivation (n=8) significantly reduces population sparseness compared to normal sleep (n=10). **(D)** Color-coded matrices displaying average pairwise correlations between Kenyon cell response patterns to multiple odors for normal sleep flies (n=10) and sleep-deprived flies (n=8). Magenta dashed boxes indicate the inter-odor correlation of similar odor pairs (4O:1M vs. 4M:1O). Orange dashed boxes indicate the inter-odor correlation of dissimilar odor pairs (4O:1M vs. PA). **(E)** Inter-odor correlation between similar odors (4O:1M vs. 4M:1O) is significantly increased in sleep-deprived flies (n=8) compared to normal sleep controls (n=10). **(F)** Inter-odor correlation between dissimilar odors (4O:1M vs. PA) remains unchanged between normal sleep controls (n=10) and sleep-deprived flies (n=8). Dots represent individual flies, lines indicate group means, and error bars represent SEM. Statistical significance was examined by Mann-Whitney test; ns, not significant; *P < 0.05.

### Sleep Loss Enhances SK Channel–Mediated Inhibition in APL Neurons

Sparse coding in the mushroom body requires a balance of excitation and inhibition, with the GABAergic anterior paired lateral (APL) neuron providing critical feedback inhibition to excitatory Kenyon cells (Figure 3A)^15,32^. Given that acute sleep deprivation disrupts sparse coding in the mushroom body, we investigated whether acute sleep loss increases the excitability of Kenyon cells, reduces the inhibitory feedback from APL neurons, or both.

**Figure 3.**
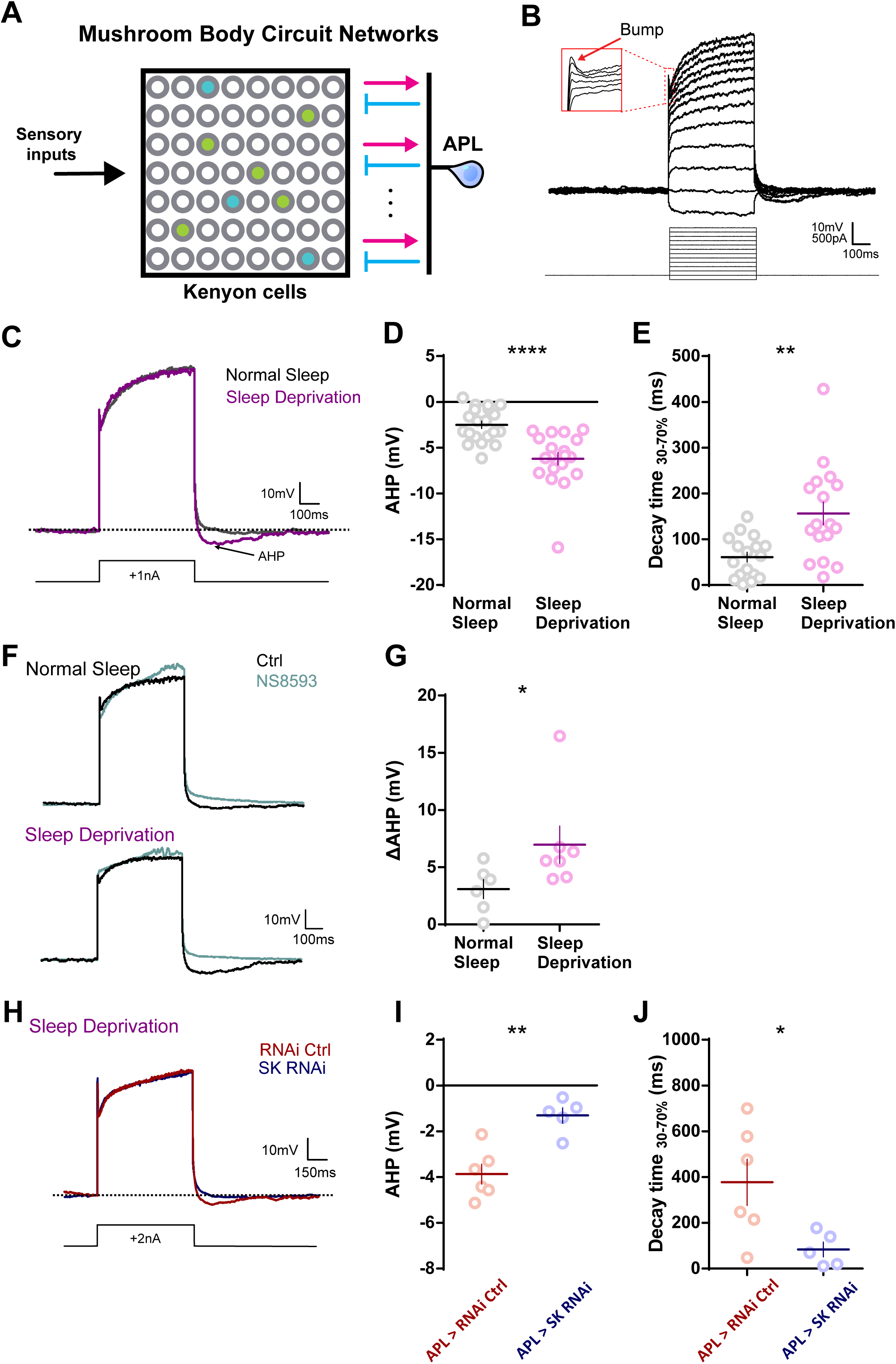
Sleep deprivation enhances SK channel-mediated afterhyperpolarization (AHP) in APL neurons in the mushroom body. **(A)** Schematic illustrating the mushroom body circuit network, highlighting the local feedback inhibition between excitatory Kenyon cells and the inhibitory anterior paired lateral (APL) neuron. **(B)** Representative current-clamp recordings show the discharge pattern of APL neurons (top) in response to current steps (bottom). The recorded APL neuron manifests a non-spiking property but exhibits initial membrane bumps upon strong depolarization (red inset). **(C)** Representative traces of AHP responses in APL neurons following 1 nA current injection after 12 hours of sleep deprivation compared to normal sleep conditions. **(D)** Quantification of AHP magnitude shows a significant increase in sleep-deprived flies (n=19) compared to those with normal sleep (n=20). **(E)** Quantification of AHP decay time (30–70%), significantly prolonged in sleep-deprived flies (n=17) compared to normal sleep flies (n=17). **(F, G)** Representative traces **(F)** and quantification **(G)** of AHP amplitude changes (ΔAHP; amplitude difference before and after treatment) induced by NS8593, a SK channel negative modulator. Sleep-deprived flies (n=7) showed significantly greater ΔAHP compared to normal sleep flies (n=6), indicating that AHP enhancement by sleep deprivation was largely SK channel-mediated. (**H–J**) Effects of APL-specific SK channel knockdown (*APL>SK RNAi*) on AHP responses following 12-hour sleep deprivation. Representative traces (**H**), AHP amplitude quantification (**I**), and AHP decay time quantification (**J**) in APL-specific SK knockdown flies (*APL>SK RNAi*, n=5) compared to RNAi control flies (*APL>RNAi Ctrl*, n=6). APL-specific SK knockdown significantly reduced AHP amplitude and decay time following sleep deprivation. Data are presented as mean ± SEM. Individual data points are shown. Statistical significance was examined by Mann-Whitney test; ns, not significant; *P < 0.05, **P < 0.01, ***P < 0.001, ****P < 0.0001.

To assess changes in excitability of Kenyon cells, we performed whole-cell patch-clamp recordings on α/β, α′/β′, and γ subtype Kenyon cells from *ex vivo* brain preparations of sleep-deprived and control (normal sleep) flies. Measurements of various membrane properties, including resting membrane potential, input resistance, afterhyperpolarization (AHP) amplitude, membrane time constant, spike threshold, and firing rate in response to current injections revealed no significant differences between the two groups (Figure S2). These findings indicate that 12 hours acute sleep deprivation does not alter the intrinsic excitability of Kenyon cells.

Next, we examined the electrophysiological properties of inhibitory APL neurons by whole-cell patch-clamp recordings (Figure S3A). Although insect APL neurons were first identified nearly two decades ago^33,34^, no whole-cell patch-clamp recordings were performed on these cells. Our recordings showed that APL neurons are non-spiking and exhibit graded responses to somatic current injection (Figure 3B). Notably, they display high leak conductance, with input resistance (∼120 MΩ; Figure S3B) nearly two orders of magnitude lower than that of Kenyon cells (∼8–11 GΩ; Figures S2D, S2K, S2R), consistent with their compartmentalized activity and localized inhibition^35^. In some cases, a spikelet-like “bump” appeared upon strong current injection (Figure 3B), suggesting the presence of voltage-gated channels despite their non-spiking nature^36,37^.

Following 12 hours sleep deprivation, APL neurons show no significant changes in resting membrane potential, input resistance, or the amplitude of bump, or threshold current for bump initiation compared to controls (Figures S3B-S3E). However, sleep-deprived APL neurons exhibited a pronounced increase in afterhyperpolarization (AHP) amplitude and prolonged AHP decay time following depolarizing current injections (Figures 3C-3E). These findings indicate that APL neurons become more hyperpolarized after activation in sleep-deprived flies, potentially reducing their inhibitory feedback to Kenyon cells and thereby compromising sparse coding in the mushroom body.

The enhanced AHP observed in APL neurons after sleep deprivation suggests an upregulation of calcium-activated potassium (K□) currents. The decay time constant of the enhanced AHP is 491.1 ± 72.17 ms (mean ± SEM), consistent with the kinetics of small conductance calcium-activated potassium channel (SK channel) mediated AHP^38^. Thus, the enhanced AHPs in APL neurons after sleep deprivation may be mediated by an increased SK channel activity. To examine this hypothesis, we applied NS8593, a selective SK channel negative modulator^39^, to *ex vivo* brain preparations from sleep-deprived flies. NS8593 abolished the AHP in both conditions (Figure 3F), and the SK-dependent AHP (NS8593 sensitive AHP) was significantly larger in sleep-deprived animals (Figure 3G). Furthermore, RNAi-mediated knockdown of SK channels specifically in APL neurons eliminated the AHP enhancement observed after sleep deprivation (Figures 3H–3J). These pharmacological and genetic experiments confirm that the increased AHP after sleep loss is mediated by SK channel.

Collectively, these results demonstrate that acute sleep deprivation enhances the conductance of SK channel in APL neurons, leading to increased AHPs, thereby prolonging their refractory period. This change occurs in the absence of alterations in Kenyon cell’s excitability or other APL biophysical properties. The enhanced AHP silences APL and reduces inhibitory feedback to excitatory Kenyon cells. This mechanism may underlie the observed disruption of sparse coding in the mushroom body and pattern separation during learning following sleep loss.

### Sleep Facilitates Pattern Separation via SK Channels in APL Neurons

To determine whether SK channels in APL neurons underlies the observed effects of sleep on pattern separation, we examine the ability of flies with SK channel knockdown specifically in APL neurons in the similar odor discrimination task with or without sleep deprivation. Control flies expressing a non-targeting RNAi exhibited normal discrimination of similar odor mixtures under baseline sleep conditions but failed to discriminate the same odors following sleep deprivation (Figures 4A and 4B), consistent with prior observations (Figures 1B and 1C). In contrast, flies with SK channel knockdown specifically in APL neurons maintained their ability to discriminate between similar odors even after sleep deprivation (Figure 4C and 4D). These results indicate that the suppression of SK channel specifically in APL neurons mitigates the detrimental effects of sleep loss on pattern separation.

**Figure 4.**
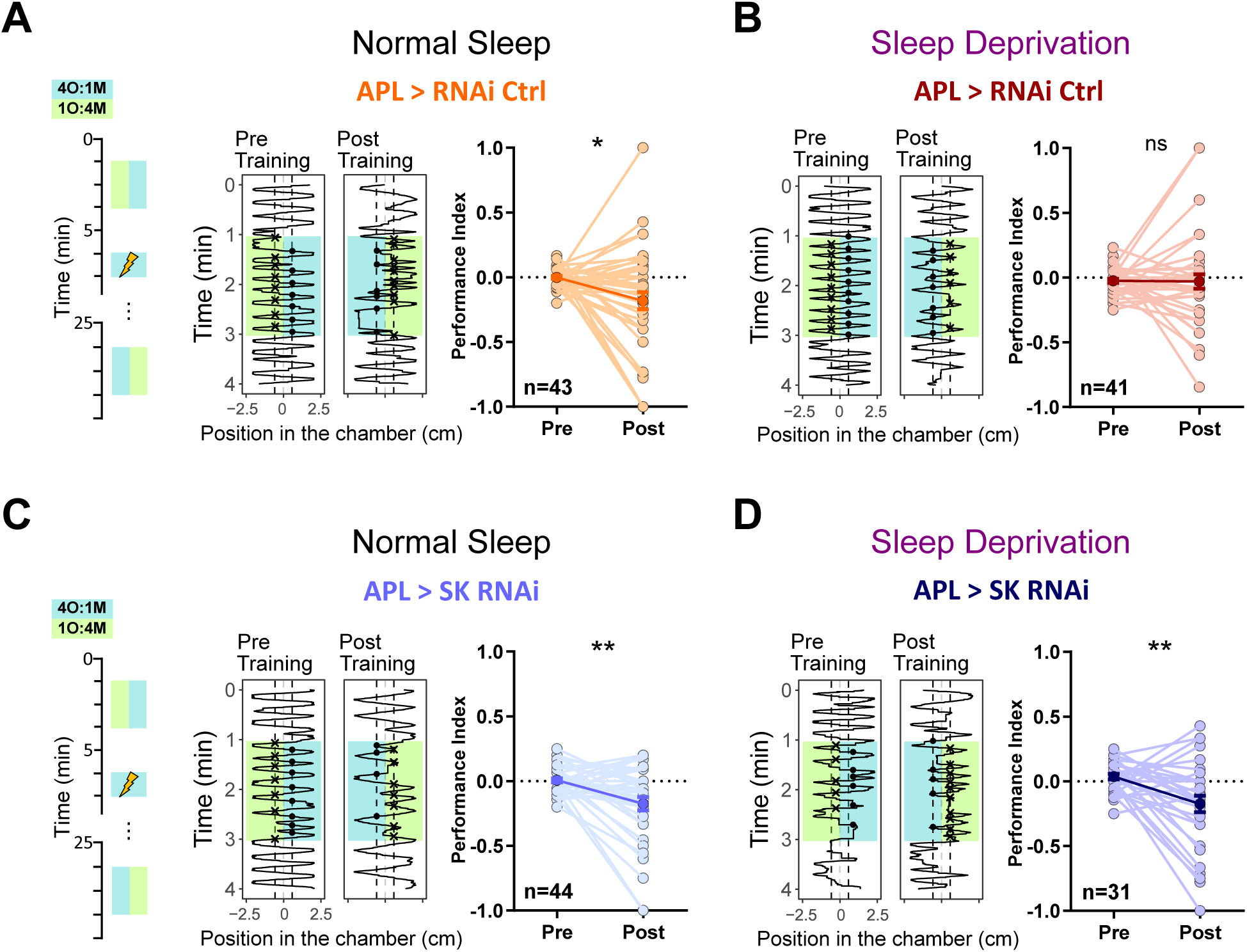
SK channel knockdown in APL neurons rescues sleep loss-induced deficits in learned discrimination of similar odors. Odor discrimination learning performance for the similar odor pair (4O:1M vs 1O:4M) was assessed in flies expressing control RNAi (*APL > RNAi Ctrl*) or SK RNAi (*APL > SK RNAi*) specifically in APL neurons under conditions of normal sleep and 12 hours sleep deprivation. **(A)** Representative trajectories (left panels) illustrate fly positions within the chamber (horizontal dimension) over time (vertical dimension). Performance indices (right panels) pre- and post-training of *APL > RNAi Ctrl* flies under normal sleep conditions (n=43) indicate successful learning discrimination. **(B)** Sleep-deprived *APL > RNAi Ctrl* flies (n=41) failed to discriminate the similar odor mixtures, as indicated by no significant difference in performance index post-training. **(C)** *APL > SK RNAi* flies under normal sleep conditions (n=31) exhibited significant learning discrimination of similar odors. **(D)** Sleep-deprived *APL > SK RNAi* flies (n=44) retained their ability to discriminate similar odors. Individual fly performance indices are shown as light-colored dots; mean values are represented by dark-colored dots. Error bars represent SEM. Statistical significance was examined by Wilcoxon matched-pairs signed rank test; ns, not significant; *P < 0.05, **P < 0.01.

## Discussion

Sleep loss impairs pattern separation, an effect observed in human studies^6,30^ but not reported in non-human animals. In this study, we found that one night of sleep deprivation impairs pattern separation in an associative learning task while leaving Pavlovian learning intact in fruit flies. This implies an evolutionarily conserved function of sleep on specific aspects of cognitive functions. Mechanistically, we discovered that sleep loss enhances the activity of small conductance calcium-activated potassium (SK) channels in GABAergic anterior paired lateral (APL) neurons, leading to increased silencing in APL neurons and reduced feedback inhibition onto excitatory Kenyon cells in the mushroom body. This reduction in feedback inhibition compromises sparse coding within the mushroom body, impairing pattern separation during associative learning. These findings highlight a mechanistic link between sleep, SK channel activity in inhibitory neurons, neural coding strategies such as sparse coding, and the preservation of cognitive functions, including pattern separation.

Our study demonstrates that sleep deprivation alters the excitability of the inhibitory APL neuron, without significantly changing excitatory Kenyon cells excitability. These observations indicate a disruption in the balance between excitation and inhibition (E/I ratio) in the mushroom body network following sleep loss. This is consistent with recent findings in the mammalian visual cortex, where feedback inhibition pathway exhibits daily oscillations and are upregulated in the morning after wakefulness in the night, whereas feedforward inhibition pathways show no significant changes^40^. Similarly, the APL neuron in the fly mushroom body network provides feedback inhibition^15,35^, suggesting that sleep’s role in maintaining feedback inhibition in neural networks is evolutionarily conserved. Although the functional consequences of sleep-dependent increases in feedback inhibition in the mammalian cortex have remained unclear, our observations provide insights into this issue. Because sparse coding is also employed in mammals, it is likely that acute sleep loss disrupts sparse coding in multiple brain areas, impairing various cognitive functions that rely on sparse coding.

SK channels are known to play a critical role in sleep regulation, particularly in maintaining low-frequency brain oscillations in mammals^41^. Studies have shown that SK2 knockout mice exhibit disrupted thalamic-cortical oscillations and fragmented sleep, underscoring their role in stabilizing sleep architecture^42^. Here, we demonstrate that sleep deprivation enhances SK channel activity in GABAergic interneurons. This suggests a bidirectional relationship in which reduced SK channel function impairs sleep, while excessive SK channel activity during wakefulness disrupts cognitive processes such as pattern separation. Understanding how sleep and wakefulness regulate SK channel activity at optimal levels in different cell types is an important question for future research.

## Methods

### Fly strains and husbandry

All *Drosophila melanogaster* strains were raised on standard cornmeal agar food at 25°C under a 12:12 h light:dark cycle with 50%-60% relative humidity. All experiments were conducted using mated female *Drosophila melanogaster*. The following stocks obtained from the Bloomington *Drosophila* Stock Center were used: *MB010B* (BDSC_68293), *UAS-SPARC2-S-mCD8::GFP* (BDSC_84148), *nSyb-IVS-phiC31* (BDSC_84152), *UAS-IVS-GCaMP6f* (BDSC_65869), *UAS-CD4-tdTomato* (BDSC_35841), *UAS-SPARC2-D-Syn21-CsChrimson::tdTomato-3.1* (BDSC_84143)*, UAS-Dcr-2.D(X)* (BDSC_24644), and *UAS-mCD8::GFP* (BDSC_29715). Stocks obtained from the Vienna *Drosophila* Resource Center for RNAi experiment were as follows: *UAS-SK RNAi*^43^ (VDRC103985), KK library control lines (*UAS-RNAi Ctrl*, VDRC60100). *LexAop-IVS-GFP-p10* is a gift from Gerald M. Rubin. *VT43924-Gal4.2*^35^ are previously generated. *VT43924-LexA::GAD* was created via LR Clonase reaction between pCR8/GW/TOPO-VT43924 and pBPnlsLexA::GADflUw^44^. The resulting construct was inserted in attP2 by the University of Cambridge Fly Facility Microinjection Service. All fly lines were outcrossed in the wild-type *w*^1118^ (BDSC_5905) background for at least 5 generations, except the lines using in the RNAi experiments. The genotype of the flies used for each experiment are summarized in Table S1.

### Sleep behavior and sleep deprivation

Flies aged 3-4 days old post-eclosion were loaded individually into the tubes containing sugar agar (2% agar with 5% sucrose) for at least one night to monitor or manipulate their sleep. Locomotion activity of individual flies was recorded using a video-based tracking device, ethoscope with an acquisition rate at 2 Hz^45^, and a sleep event was defined by the inactive state for 5 min. Mechanical rotation of individual tubes for 1.5 s was triggered by immobility of respective flies for consecutive 10 s to achieve sleep deprivation with a closed-loop system as previously used in the lab^46^. To exclude the outliers from sleep and sleep-deprived flies, a cutoff was set to 40% of sleep time in the previous night, yielding two non-overlapping groups with a Cohen’s *d* of 1.89. The successful rate of sleep deprivation in this study is >95%. For both sleep and sleep-deprived conditions, experiments were acquired at ZT0-3 with or without sleep deprivation one night before. For the condition of sleep recovery, experiments were conducted at ZT3-6. Sleep analysis was performed with adaptations to the rethomics pipeline^47^ in R.

### Learning behavior of individual flies

Olfactory learning behavior of individual flies was conducted in a versatile custom-built apparatus consisting of 20 chambers with two air inlets at each side and two vents in the middle of individual chambers^29^. Chambers consisted of 3D-printed enclosures (CPE, Ultimaker) with two glass slides as floors and ceilings, where the dimension of a chamber (in mm) was 50 × 5 × 1.5 (l × w × h). Chambers equipped with transparent glass slides coated with ITO (indium tin oxide) were applied to deliver electric shocks during aversive conditioning, whereas chambers equipped with non-ITO transparent glass slides were used in post-conditioning preference tests to avoid contextual memory confounding the performance in olfactory discrimination tasks. ITO coating with a grid pattern on the glass slides was designed to insulate the positive and negative electrodes (a hundred ITO electrodes with a width of 0.5 mm spaced by 0.1 mm apart). ITO glasses were connected to a 75-V DC source through relay modules to control delivery of electrical shocks. Chambers were illuminated by 940 nm LEDs below and recorded by a top camera (Basler ace) equipped with a Computar M0814-MP2 lens at a frame rate of 5 Hz. Real-time activity of individual flies was imaged using computerized video-tracking. Filtered and humidified air was tuned by a main flow controller and was split into two streams (2 l/min for each) for either side. The stream of each side was independently controlled by solenoid valves to switch the flow between the vials containing mineral oil or diluted odors in mineral oil and further distributed evenly to 20 chambers at a flow rate of 0.1 l/min for each side. Odors were diluted in the mineral oil at the following concentration: 1:375 for MCH (4-methyl-cyclohexanol, CAS No. 589-91-3, Sigma-Aldrich), 1:500 for OCT (3-octanol, CAS No. 589-98-0, Sigma-Aldrich), 1:10000 for PA (pentyl acetate, CAS No. 628-63-7, Sigma-Aldrich). Odor blends were mixed via manipulating the proportion of the air flow through the vials in which two sub-streams flowed at the rate of 0.4 l/min and 1.6 l/min for MCH and OCT, or vice versa. Odors were vented from the middle of chambers to a collection bottle for absorption of excessive odor.

Odors were introduced to the chambers for 2 min to access the preference before and 20 min after aversive conditioning; only flies with performance index (PI) ranging from −0.25 to 0.25 in the preference test before conditioning were included. For aversive conditioning, flies were exposed to CS^+^ odor for 1 min paired with twelve electric shocks at 0.2 Hz. 14 mm in the middle of chambers was defined as a decision zone, and the incidences entering from decision zone toward either side during the delivery of odors were quantified to calculate the PI of individual flies by the following equation:

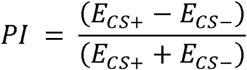

Where E*_CS+_* represents the number of entrances into zone with CS^+^ odor and E*_CS-_* represents the number of entrances into zone with CS^-^ odor or air.

### *In vivo* Calcium imaging

Flies were briefly cold-anesthetized and secured to a custom imaging holder by threading a fine string beneath the neck to immobilize the head. Wings, eyes, and the dorsal thorax were fixed using UV-curable glue, and the proboscis and legs were immobilized to minimize motion artifacts and suppress dopaminergic modulation associated with voluntary leg movements. A small window of cuticle was removed to expose the brain, and overlying air sacs and fat bodies on the posterior surface were gently cleared. All surgical procedures were performed in external saline containing (in mM): 103 NaCl, 3 KCl, 1.5 CaCl□, 4 MgCl□, 26 NaHCO□, 5 TES, 1 NaH□PO□, 10 trehalose, and 10 glucose (pH 7.3, bubbled with 95% O□/5% CO□, 275 mOsm). Following dissection, the fly was transferred to a recording stage under a two-photon microscope (FemtoSmart Dual, Femtonics) and continuously superfused with the same saline. GCaMP fluorescence was excited using a mode-locked femtosecond laser (Chameleon Ultra II, Coherent) tuned to 920 nm, and emission was detected with a GaAsP detector (Hamamatsu Photonics). For each fly, a single imaging plane was selected to include ∼200 clearly identifiable Kenyon cells, and a 3-min rest period was allowed before image acquisition. Images were acquired at 5 Hz using a galvo scanner and a 20× water-immersion objective (XLUMPLFLN20XW, Olympus), with a pixel resolution of 0.47 × 0.47 μm. Image acquisition and microscope control were performed using MESc v3.5 (Femtonics), and image series were exported as TIFFs for further analysis.

### Odor stimulation

Odors were diluted in MO (mineral oil, CAS No. 8042-47-5, Sigma-Aldrich) at the following concentrations: 1:37.5 for MCH (4-methyl-cyclohexanol, CAS No. 589-91-3, Sigma-Aldrich), 1:50 for OCT (3-octanol, CAS No. 589-98-0, Sigma-Aldrich), 1:1000 for AAC (Acetic acid, CAS No. 64-19-7, Honeywell), 1:1000 for PA (pentyl acetate, CAS No. 628-63-7, Sigma-Aldrich). For two mixed odor blends, 4M:1O and 4O:1M were 1:46.875 for MCH and 1:250 for OCT and 1: 187.5 for MCH with 1:62.5 for OCT, respectively. Filtered and humidified mainstream (1.8 l/min) air was mixed with a sub-stream (0.2 l/min) flowed through vials filled with 10ml mineral oil, and the sub-stream was directed to other vials containing diluted odorant in 10 ml mineral oil via a custom-built solenoid valve system while receiving a TTL pulse from the microscope. The air/odor streams were split into an exhausted stream and a flow-controlled stream to the delivery tube with a flow rate of 0.15 l/min, which shorten the delay time to 1s from odor vials to the end of delivery tube and minimize the onset and offset artifact when the solenoid valve switches. Every fly received two series of odor presentation trials in a pseudo-random sequence in which the same odor was not given consecutively. Among all olfactory stimuli, mineral oil was included as a blank stimulus, it was always presented first in every recording. Each odor was given twice to ensure PNs processed the olfactory stimuli faithfully^48^, so only the response to each odor in the second time was adopted for data analysis. Imaging was incessantly acquired for 420 s over 14 odor presentation trials in which every trial consists of a 250 ms odor pulse triggered 5 s after the trial onset and with an inter-stimulus interval for 30 s.

### Image processing and analysis

Fluorescence image stacks were processed and analyzed using ImageJ/Fiji software. Motion correction was first applied to all imaging sessions using the Fiji plugin TurboReg to correct the drift in the x-y plane. Recording with significant z-drift, uncorrectable motion artifacts, or obvious structure distortion were excluded from further analysis. After image registration, regions of interest (ROIs) corresponding to individual Kenyon cell somata in the mushroom body were manually identified from the motion-corrected average intensity projection image across all frames. Using ImageJ’s ROI Manager, circular ROIs (typically 3–5 µm in diameter) were drawn around clearly distinguishable somatic boundaries based on baseline fluorescence morphology. Only ROIs with consistent shape, size, and signal quality across trials were retained. Approximately 150-200 Kenyon cells were selected per preparation depending on visibility. For each ROI, fluorescence intensity was extracted over time and used to calculate the fractional change in fluorescence (Δ*F/F_0_*). The baseline fluorescence (*F_0_*) was the mean intensity of 6 s window before odor was delivered to the fly. Peak response (*F_t_*) is calculated as the average fluorescence of 6s window after baseline. Within each trial, Δ*F* was calculated as the difference between peak and baseline fluorescence, and Δ*F/F_0_* was then calculated as the Δ*F* divided by the baseline fluorescence (*F_0_*) shown as the following equation:

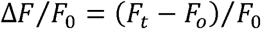

### Population sparseness

To access the sparseness of odor representations in the mushroom body, olfactory responses to each stimulus across multiple trials were computed for each ROI. Response of the ROI was significant if the peak response was 3 s.d. (σ) greater than the baseline, otherwise the ROI was considered unresponsive (A 3σ threshold corresponds to the top ∼0.3% of a normal distribution.) Population sparseness (Sp) was calculated for each odor across all ROIs (e.g., Kenyon cells) using the following equation^20^:

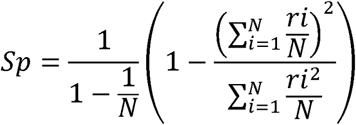

Where ri is the normalized response (Δ*F/F_0_*) of ROIi to the odor and N is the total number of ROIs. For those responses

### Inter-Odor correlation

To assess the similarity of population representations across odors, we computed the Pearson correlation coefficient between the response vectors (across all ROIs) for each pair of odors. For a given pair of odors A and B:

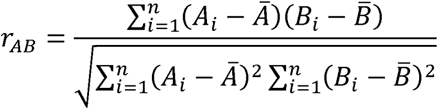

Where A_i_ and B_i_ are responses of ROI_i_ to odors A and B, respectively. Correlation matrices were visualized to compare representational overlaps across the odor panel.

### Electrophysiological recordings

Flies were anesthetized on ice shortly before decapitation. Brains were dissected in the same external saline used for the calcium imaging experiments (see above), and the glial sheath surrounding the brain was removed to visualize the recording site. The dissected brain was immobilized on the recording chamber using a custom-made platinum anchor. Cell somata were targeted for patching under a 20× water-immersion objective (XLUMPlanFL, Olympus) and visualized using IR-DIC on an upright BX51WI microscope (Olympus) with the Slicescope (Scientifica). Before patching, the perineural sheath was further removed and to expose a clean surface using a suction pipette (∼1 MΩ) connected to the pipette holder. Patch pipettes were fabricated from borosilicate glass capillaries (1.5 mm o.d./ 0.84 mm i.d., 1B150F-4, World Precision Instruments) using a Sutter P-1000 puller (Sutter Instrument). For Kenyon cell recordings, the pipettes were fire-polished by a microforge (MF-830, Narishige) under a ∼35 psi air pressure backfilled by a picospritzer (Harvard Apparatus) to achieve a final resistance of 11–14 MΩ^49,50^. For APL recordings, the pipettes were fire-polished without a backfilled pressure to achieve a typical resistance of 6–9 MΩ. Polished pipettes were filled with the internal solution containing (in mM): 140 potassium gluconate, 10 HEPES, 1 EGTA, 4 MgATP, 0.5 Na GTP, 1 KCl, 13 biocytin hydrazide. Whole-cell recordings were made with a MultiClamp 700B amplifier (Molecular Devices) equipped with a CV-7B headstage, lowpass-filtered at 3 kHz, and sampled at 10 kHz using a Digidata 1550B digitizer controlled through pCLAMP 10 (Molecular Devices). Data were corrected for liquid junction potential (Neher, 1992), and data analysis was performed in Clampfit (Molecular devices). Putative Kenyon cells were selected based on the anatomical location without knowing the cell types during recording, and only recordings with recognizable morphology of Kenyon cells were included. Only one recording was tempted from one hemisphere for a clean post-hoc identification of Kenyon cell types. APL recordings were guided by visualization of green fluorescence protein under epifluorescence, and partial tissue covering on the APL soma were removed if necessary. Resting membrane potential was taken immediately after break-in without holding current and compensation for the pipette capacitance and series resistance. After that, all properties (Figures 3 and S2) were recorded with the neutralization of pipette capacitance and bridge balance for series resistance in current clamp. Cells were held at around −60 ± 5 mV by injecting a hyperpolarizing current.

For Kenyon cell recordings, a 600 ms current ramp, from –2 to +30 pA, starting at a membrane potential of –60 ± 5 mV was injected to estimate spike threshold and afterhyperpolarization (AHP). The spike onset was detected by finding maximum in the time derivative of membrane potential in the ramp trace in which each spike waveform should contain clear upward and downward deflections for the distinction of spikes from EPSPs, and the membrane potential corresponding to the first spike onset in the ramp was defined as the spike threshold. AHP was detected by the downward peak following the first spike and was estimated by the difference between spike threshold and downward peak. Input resistance, membrane time constant, and firing rate were estimated by a series of current steps consisting of a 500 ms hyperpolarizing current (–1 pA) followed by 750 ms current steps (1 pA increments, starting at −5 pA) 1 s apart from the constant hyperpolarizing step. Input resistance was calculated from the average change of steady-state voltage elicited by the constant hyperpolarizing current. Membrane time constant was determined by fitting a single exponential to the averaged voltage deflection responded to the constant hyperpolarizing current. Firing rate was the spike counts elicited by the 750ms current steps. After the recording, the brain was immediately fixed in 4% paraformaldehyde at room temperature for 20 min for further post-hoc staining.

For APL recordings, input resistance, membrane time constant, AHP were estimated by a series of current steps consisting of a 500 ms hyperpolarizing current (–50 pA) followed by 500 ms current steps (50 pA increments, starting at −100 pA) 1 s apart from the constant hyperpolarizing step. Bump responses and AHPs were elicited by high discharge in membrane potential, so the amplitudes were estimated by the deflection of membrane potential in response to 1 nA current injection during charging and after deactivation, respectively. To assess the channel contribution to the enhanced AHP in sleep-deprived flies, we estimated the decay time constant by exponential curve fitting. However, this method is not applicable in the normal sleep group, since many recordings exhibited low-amplitude AHPs, for which exponential fitting yielded unreliable decay estimates. Therefore, for cross-condition comparisons, AHP kinetics were quantified as the time required for the membrane potential to decay from 70% to 30% of the peak AHP amplitude. For pharmacological experiments, NS8593 (492031, Sigma-Aldrich), rather than another selective inhibitor - apamin, was applied to block *Drosophila* SK channels, since fly SK has been reported insensitive to apamin^51^. 10 mM stock of NS8593 in dimethyl sulfoxide (DMSO, CAS No. 67-68-5, Sigma-Aldrich) was dissolved in the external saline, yielding NS8593 at a final concentration of 10 uM with 0.1 % DMSO (v/v). 0.1% DMSO (v/v) was added in the external saline as vehicle control, and the difference on AHP amplitude in response to 1 nA current was calculated by the responses before and 20 min after treatment of NS8593.

### Immunofluorescence and confocal imaging

Fly brains acquired after electrophysiology experiments were washed in PBS with 0.5% Triton X-100 (PBST) for 15 min each time. After washing for three times in PBST, brains were incubated in PBST containing 5% normal goat serum for 25 min at room temperature, and then incubated with mouse nc82 (Development Studies Hybridoma Bank, 1:50) overnight at 4 °C. Subsequently, brains were incubated with secondary antibodies including Alexa 488 or Alexa 647 anti-mouse (Invitrogen, 1:400) and Alexa 488- or Alexa 647-conjugated streptavidin (Invitrogen, 1:400) for 2 hours at room temperature. Brains were washed in PBST three times for 15min at room temperature after every incubation step with antibodies or Alexa Fluor Dyes. After the final washing step, brains were mounted in Vectashield (Vector Labs), and images were acquired using Nikon NiE A1R confocal microscopy with a 1µm step-size under 40x magnification.

### Statistical analyses

Data were analyzed using Prism 10 (GraphPad). For comparisons of two matched groups, paired t-test or Wilcoxon’s matched pairs test were applied for normally or non-normally distributed data, respectively. For comparison of unpaired groups, unpaired t-test or Mann-Whitney U-tests. Current versus firing rate relationships were analyzed using Repeated measures two-way ANOVA, followed by Bonferoni post-hoc tests.

## Acknowledgments

We thank the Bloomington Drosophila Stock Center, Dr. Gerald M. Rubin for providing the fly stocks. This research was supported by the European Research Council (StG 758580 to S.L.), Fonds voor Wetenschappelijk Onderzoek (FWO) Research Project (G052922N to S.L.).

