## Supplemental Figures for "Sleep Facilitates Pattern Separation through SK Channel-Mediated Sparse Coding"

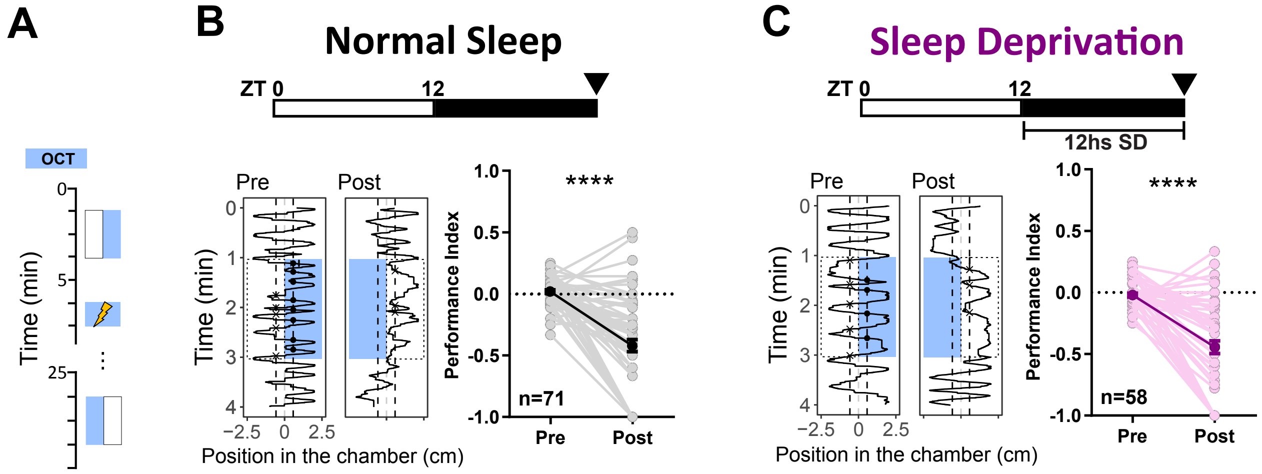


**Figure S1. Acute sleep deprivation does not impair learning performance in an aversive olfactory conditioning task.** (**A**) Experimental paradigm illustrating the aversive olfactory conditioning protocol, comprising pre-training test, aversive conditioning (training), and post-training test phases plotted vertically against time. Colored rectangles indicate the timing of odor (OCT) presentations in the chamber.

(**B** and **C**) Learning performance of flies under conditions of normal sleep (**B**, n=71) or following 12 hours of sleep deprivation (**C**, n=58). Top panels: Bars indicate the 12:12 light-dark cycles (white: light, black: dark); Zeitgeber Time (ZT) 0 marks lights-on; arrows indicate the timing of behavioral testing. Bottom-left panels: Representative trajectories of individual fly positions in the testing chamber (horizontal axis: chamber position, vertical axis: time) before (Pre) and after (Post) aversive conditioning. Bottom-right panels: Quantification of learning, represented by the performance index. Both normal sleep and sleep-deprived (SD) flies exhibited robust learning, with significantly reduced performance indices post-conditioning.

Individual fly performance indices are shown as light-colored dots; mean values are indicated by dark-colored dots. Error bars represent SEM. Statistical significance was examined by Wilcoxon matched-pairs signed rank test; ****$P < 0.0001$.

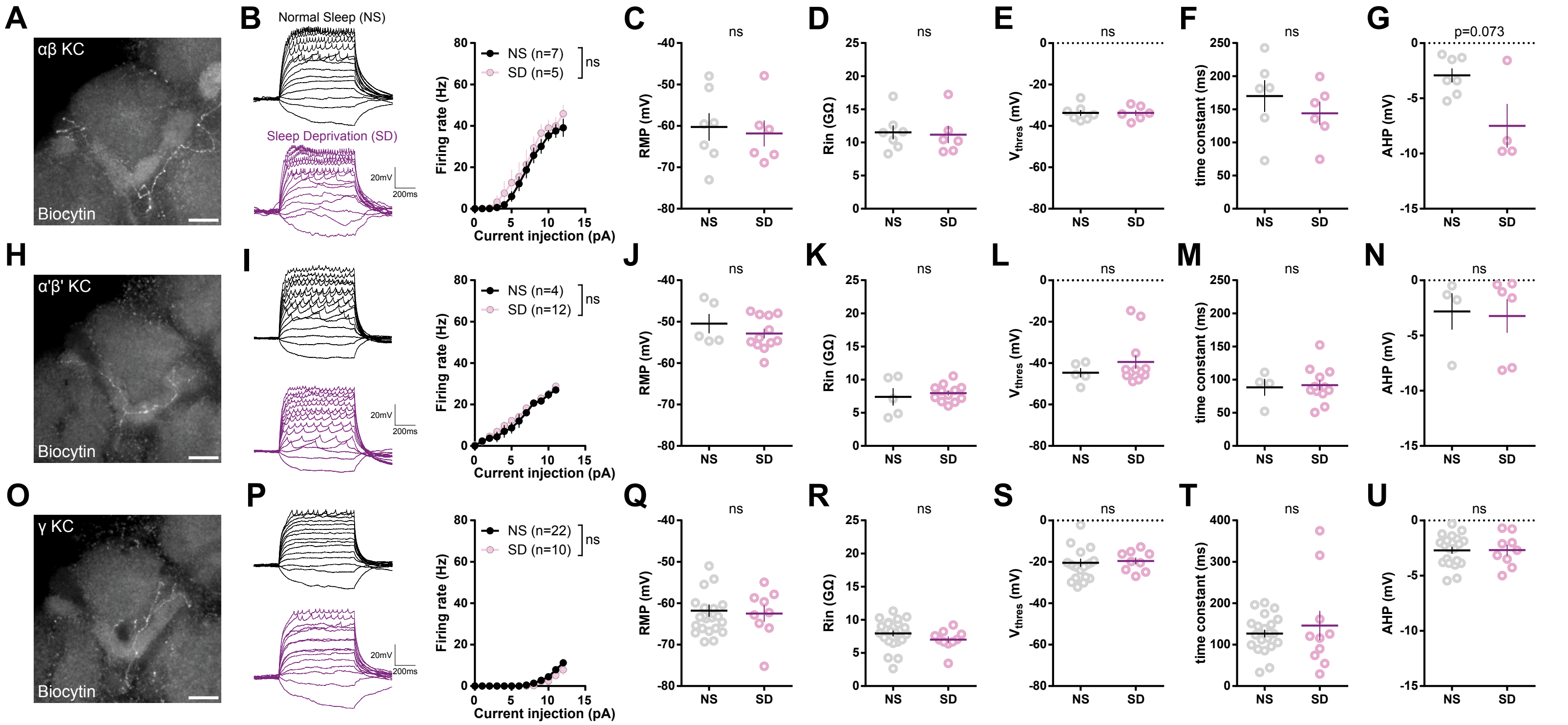


**Figure S2. Sleep deprivation does not alter the intrinsic electrical properties of Kenyon cells (KCs).**
(**A–G**) Intrinsic membrane properties of αβ KCs under normal sleep (NS) and sleep deprivation (SD) conditions.
(**A**) Post-hoc immunohistochemistry of a representative biocytin-filled αβ KC. Scale bar, 20 $\mu$m. (**B**) Left: representative voltage traces from αβ KCs in NS (black) and SD (magenta) flies in response to current steps (−1 to 12 pA).
Right: current–firing rate relationships show no significant difference between groups. (**C–G**) Quantification of resting membrane potential (RMP; **C**), input resistance ($R_{in}$; **D**), spike threshold ($V_{thres}$; **E**), membrane time constant (**F**), and afterhyperpolarization (AHP) amplitude (**G**) reveals no significant differences, though a trend toward larger AHP in SD flies is noted ($p = 0.073$).
(**H–N**) Intrinsic membrane properties of α′β′ KCs under NS and SD conditions. (**H**) Post-hoc immunohistochemistry of a representative biocytin-filled α′β′ KC. Scale bar, 20 $\mu$m. (**I**) Left: representative firing responses to current steps in NS and SD flies; right: no significant difference in current–firing rate curves. (**J–N**) No significant differences were observed in RMP (**J**), $R_{in}$ (**K**), $V_{thres}$ (**L**), membrane time constant (**M**), or AHP amplitude (**N**).
(**O–U**) Intrinsic membrane properties of γ KCs under NS and SD conditions. (**O**) Post-hoc immunohistochemistry of a representative biocytin-filled γ KC. Scale bar, 20 $\mu$m. (**P**) Left: representative voltage responses to current injections; right: no significant difference in firing rate across conditions. (**Q–U**) RMP (**Q**), $R_{in}$ (**R**), $V_{thres}$ (**S**), membrane time constant (**T**), and AHP amplitude (**U**) were not significantly different between NS and SD groups.
Data are presented as mean ± SEM. Individual data points are shown. Statistical comparisons were performed using Mann–Whitney test. ns, not statistically significant.

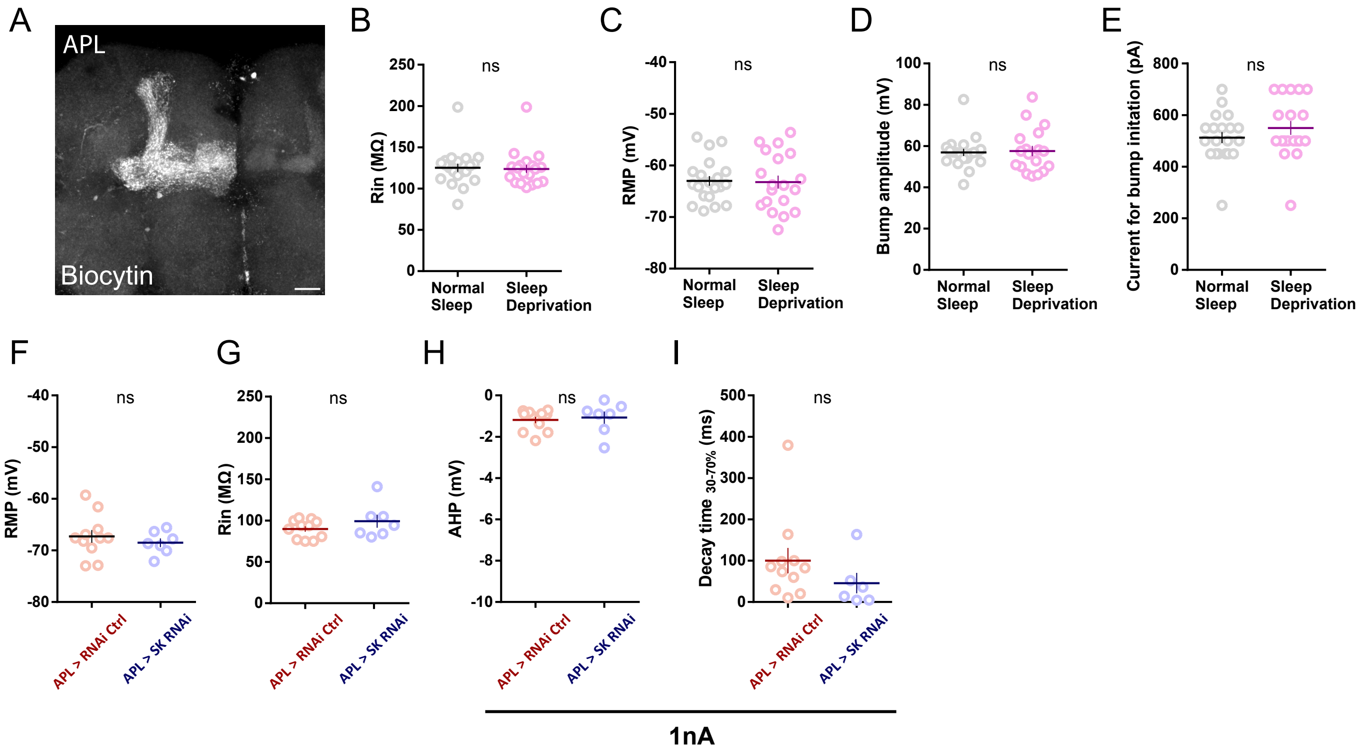


**Figure S3. Intrinsic membrane properties of APL neurons are unaffected by sleep deprivation or genetic knockdown of SK channels.**
(**A**) Post-hoc immunohistochemistry of an APL neuron filled with biocytin during whole-cell recording. Scale bar, 20 $\mu$m.
(**B–E**) Quantification of intrinsic membrane properties of APL neurons from normal sleep (NS) and sleep-deprived (SD) flies shows no significant differences in (**B**) input resistance ($R_{in}$; NS, NS, n=20 ; SD, n=19), (**C**) resting membrane potential (RMP; NS, n=20 ; SD, n=19), (**D**) maximal amplitude of the initial bump response (NS, n=20 ; SD, n=19), (E) threshold current required to evoke a bump (NS, n=20 ; SD, n=19).
(**F–I**) Intrinsic properties of APL neurons recorded from SD flies expressing control RNAi (*APL > RNAi Ctrl*) or SK RNAi (*APL > SK RNAi*). No significant differences were observed between genotypes in (**F**) RMP (RNAi Ctrl, n=11; SK RNAi, n=7) and (**G**) $R_{in}$ (RNAi Ctrl, n=11; SK RNAi, n=7). (H) AHP amplitude was comparable between genotypes in response to a 1 nA current injection after 12 hours of sleep deprivation. (RNAi Ctrl, n=11; SK RNAi, n=7) (**I**) No significant differences were found in AHP decay time in response to a 1 nA current injection. (RNAi Ctrl, n=11; SK RNAi, n=7)
Data are presented as mean ± SEM. Individual data points are shown. Statistical significance: ns, not significant.
