## Supplemental Table 1 for "Sleep Facilitates Pattern Separation through SK Channel-Mediated Sparse Coding"

| Figure | Genotypes |
| --- | --- |
| 1 | +; nsyb-phiC31, R13F02-p65.AD/ UAS-SPARC2-S-mCD8::GFP; R52H09-GAL4.DBD/+ |
| 2 | +; R13F02-p65.AD/ UAS-IVS-GCaMP6f, UAS-CD4-tdTomato; R52H09-GAL4.DBD/+ |
| 3B-G | +; nsyb-phiC31, R13F02-p65.AD/ UAS-SPARC2-D-Syn21-CsChrimson::tdTomato-3.1; R52H09-GAL4.DBD/ VT43924-LexA::GAD, LexAop2-IVS-myr::GFP |
| 3H-J | UAS-Dcr-2.D/+; UAS-mCD8::GFP/ UAS-dSK RNAi; VT43924-Gal4.2/+ |
| 4 | UAS-Dcr-2.D/+; UAS-mCD8::GFP/ UAS-dSK RNAi; VT43924-Gal4.2/+ |
| S1 | +; nsyb-phiC31, R13F02-p65.AD/ UAS-SPARC2-S-mCD8::GFP; R52H09-GAL4.DBD/+ |
| S2 | +; nsyb-phiC31, R13F02-p65.AD/ UAS-SPARC2-D-Syn21-CsChrimson::tdTomato-3.1; R52H09-GAL4.DBD/+ |
| S3B-E | +; nsyb-phiC31, R13F02-p65.AD/ UAS-SPARC2-D-Syn21-CsChrimson::tdTomato-3.1; R52H09-GAL4.DBD/ VT43924-LexA::GAD, LexAop2-IVS-myr::GFP |
| S3F-I | UAS-Dcr-2.D/+; UAS-mCD8::GFP/ UAS-SK RNAi; VT43924-Gal4.2/+ |
